# Onset of infectiousness explains differences in transmissibility across *Mycobacterium tuberculosis* lineages

**DOI:** 10.1101/2024.08.27.609909

**Authors:** Etthel M. Windels, Cecilia Valenzuela Agüí, Bouke C. de Jong, Conor J. Meehan, Chloé Loiseau, Galo A. Goig, Michaela Zwyer, Sonia Borrell, Daniela Brites, Sebastien Gagneux, Tanja Stadler

## Abstract

*Mycobacterium tuberculosis* complex (MTBC) lineages show substantial variability in virulence, but the epidemiological consequences of this variability have not been studied in detail. Here, we aimed for a lineage-specific epidemiological characterization by applying phylodynamic models to genomic data from different countries, representing the most abundant MTBC lineages. Our results show that all lineages are associated with similar durations and levels of infectiousness, resulting in similar reproductive numbers. However, L1 and L6 are associated with a delayed onset of infectiousness, leading to longer periods between subsequent transmission events. Together, our findings highlight the role of MTBC phylogenetic diversity in tuberculosis disease progression and transmission.

## Introduction

Human tuberculosis (TB) is characterized by a large heterogeneity in clinical and epidemiological features (Coscolla and Gagneux 2010; Cadena et al. 2017). Although several host and environmental factors partially underlie this variability, there is increasing evidence that genetic diversity within the *M. tuberculosis* complex (MTBC) also plays a role in the disease presentation and transmission dynamics. Ten human-adapted MTBC lineages (L1 to L10) have been identified to date (reviewed in Orgeur et al. 2024; Guyeux et al. 2024). L1 to L6 are globally the most abundant lineages, where L1, L5, and L6 are often called the “ancient” lineages, and L2, L3, and L4 are commonly referred to as the “modern” lineages (Brosch et al. 2002; Gagneux 2018; Bottai et al. 2020).

Several animal and macrophage infection studies have shown reduced virulence of strains belonging to “ancient” lineages compared to strains from “modern” lineages, observed as restricted *in vivo* growth, host immune modulation, and disease severity (reviewed in Coscolla and Gagneux 2010; Coscolla and Gagneux 2014; Tientcheu et al. 2017; Peters et al. 2020). Consistent with this reduced virulence, molecular epidemiological studies have reported a lower transmissibility of “ancient” compared to “modern” lineages (Yang et al. 2012; Guerra-Assunção et al. 2015; Asare et al. 2018; Holt et al. 2018; Sobkowiak et al. 2020; Freschi et al. 2021; Zwyer et al. 2023; Gröschel et al. 2024), with the majority of these studies using clustering rates and/or terminal branch lengths (TBLs) to quantify transmission. These metrics indirectly estimate the time between subsequent transmission events, but do not explicitly consider patient infectiousness (i.e. the ability to transmit) during that time, resulting in an incomplete picture of the transmission dynamics for two reasons. First, the time between transmission events also includes the time between infection and the onset of infectiousness, and is hence not necessarily a measure for how rapidly infectious patients spread the disease. Second, the duration of infectiousness directly affects the effective reproductive number (R_e_), i.e. the expected number of secondary cases caused by a single infected individual, which determines whether case numbers increase or decrease over time. An additional limitation of clustering rates and TBLs is that they ignore (potentially lineage-specific) variation in clock rate and sampling intensity, which both affect the genetic distance between sampled isolates (Menardo 2022).

Phylodynamic birth-death models explicitly model the processes of molecular evolution, transmission, becoming (non-)infectious, and sampling (Stadler 2009; Stadler 2010; Kühnert et al. 2016), which allows disentangling the relevant evolutionary and epidemiological characteristics. Here, we used different versions of the birth-death model to characterize the epidemiological features of the main MTBC lineages in unprecedented detail. As we aimed for a comprehensive epidemiological comparison of lineages, the models were applied to genomic data from four different sampling locations where “ancient” (L1 and L6) and “modern” (L2, L3, and L4) lineages co-circulate (Malawi, Tanzania, The Gambia, and Vietnam). Our results suggest that all lineages (except for L1 in Vietnam) are characterized by an R_e_ close to 1, and are therefore on average not expected to change much in relative abundance over time. However, L1 and L6 are characterized by significantly longer periods between subsequent infection events than lineages L2, L3, and L4. We show that this is due to a delayed onset of infectiousness in L1 and L6-infected individuals, rather than an overall reduced level of infectiousness. These findings provide new insights into the implications of MTBC genetic diversity on transmission and disease progression.

## Results

We used publicly available whole-genome sequencing data from four countries where “ancient” and “modern” lineages co-circulate: 1,684 sequences from Malawi (L1, L2, L3, and L4) (Guerra-Assunção et al. 2015), 921 sequences from Tanzania (L1, L2, L3, and L4) (Zwyer et al. 2023), 1,086 sequences from The Gambia (L2, L4, and L6) (Gehre et al.), and 1,623 sequences from Vietnam (L1, L2, and L4) (Holt et al. 2018) (see Materials and Methods for details on the study populations).

By applying phylodynamic birth-death models to these genomic data, we aimed for a detailed epidemiological characterization of the different MTBC lineages, including their associated onset and duration of infectiousness, level of infectiousness (rate of secondary transmissions per infected individual), and the resulting reproductive number. As the classical birth-death model (also called “single-type birth-death model”) does not distinguish between non-infectious and infectious infected individuals, we used an extension based on the multi-type birth-death model, allowing for an initial non-infectious period in infected individuals (Figure 1). This is achieved by introducing an epidemiological compartment representing infected non-infectious individuals in the population. The non-infectious/infectious dichotomization in this model is purely based on the ability to cause secondary infections, rather than on symptoms or radiographic/microbiological markers. This simplifies the continuous clinical disease spectrum characterizing TB infection and disease (Drain et al. 2018; Kendall et al. 2021), but potentially represents a major improvement in accuracy compared to the single-type birth-death model, as well as less fine-grained transmissibility metrics. This multi-type birth-death model was fitted to the genomic data from Malawi, Tanzania, The Gambia, and Vietnam, with priors reported in Table S1 (see Materials and Methods for details on the model). Except for L1 in Vietnam, the posterior estimates for the R_e_ were close to 1 for all lineages in all sampling locations (Figure 2a). This suggests that in these countries, the relative abundance of each lineage is on average not changing considerably over time. However, the average time between infection and onset of infectiousness was consistently estimated to be longest for L1 and L6 (Figure 2b). These posterior estimates are estimates of the population average, and can be translated into population distributions of times until onset of infectiousness (Figure 2c), showing that on average, 26% (L1), 73% (L2), 54% (L3), 43% (L4), and 23% (L6) of infected individuals were estimated to progress to infectious TB within one year of infection (Table 1). In contrast to this lineage-specific initial non-infectious period, the subsequent infectious period and the transmission rate (i.e. the rate of secondary case generation per infected individual) during this infectious period did not show significant differences between lineages (Figure 2d-e). Together, these results suggest that all lineages are characterized by a similar duration and level of infectiousness, but that “ancient” lineages L1 and L6 are associated with a longer period of initial non-infectiousness, and hence longer periods between infections and secondary transmission events.

**Figure 1:**
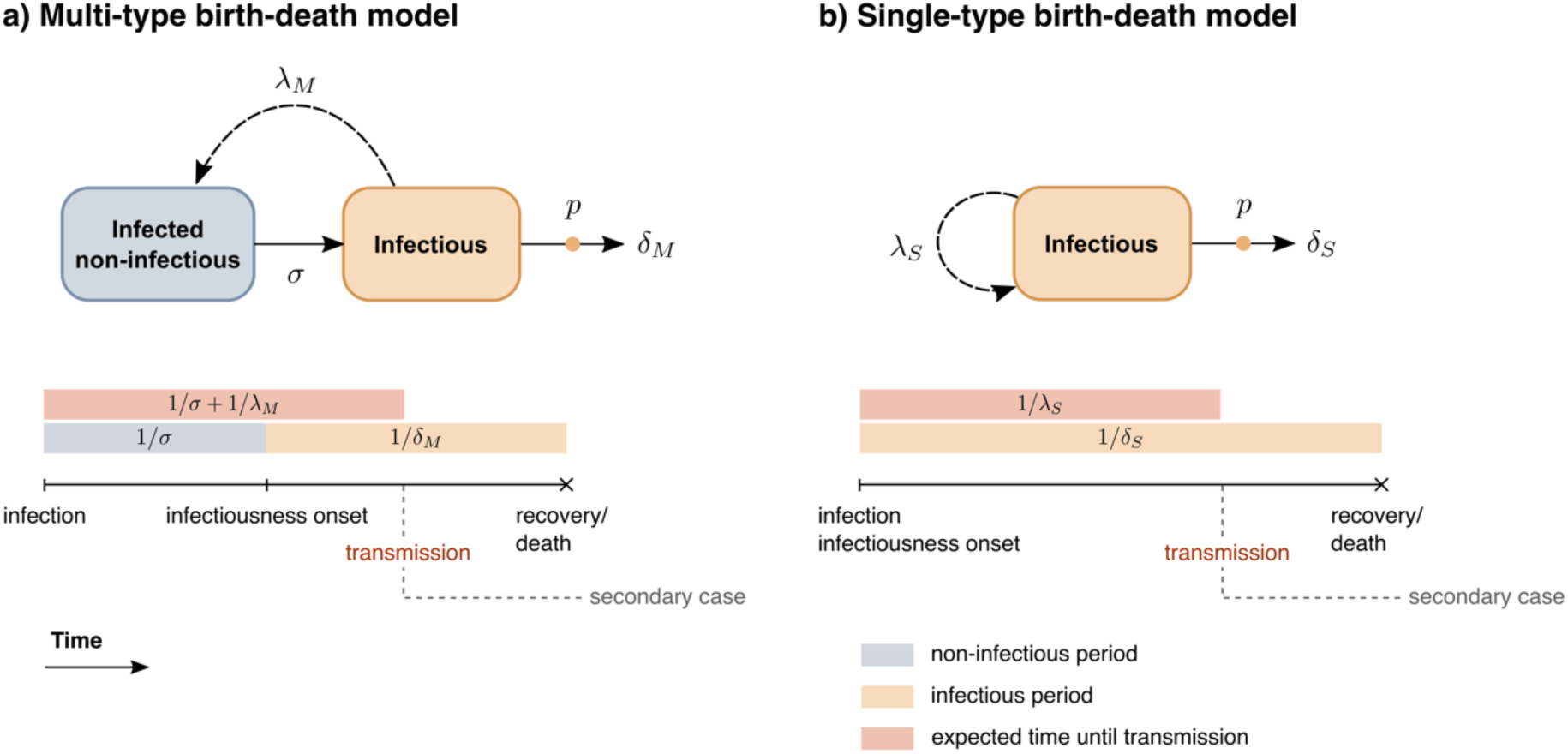
Schematic representation of the phylodynamic birth-death models used in this study. a) The multi-type birth-death model distinguishes individuals who are infected but not yet infectious from individuals who are infectious and generate secondary infections at rate λ_M_. A new infection results in a new individual in the “infected non-infectious” compartment, who moves to the “infectious” compartment with rate σ. Hence, the expected duration of the non-infectious period equals 1/σ. Due to this initial non-infectious period, 1/λ_M_ represents the expected time until secondary infection since the start of the infectious period. Individuals become non-infectious through recovery or death at rate δ_M_, resulting in an average infectious period of 1/δ_M_. b) The classical, single-type birth-death model is a more constrained version of a) including only a single compartment of infected individuals. Upon infection, individuals instantaneously become infectious, with a constant transmission rate λ_S_. Hence, the expected time between infection and the first transmission event equals 1/λ_S_. Individuals become non-infectious through recovery or death at rate δ_S_. Consequently, the average infectious period in this model equals 1/δ_S_ and corresponds to the total duration of infection. In both models, patients are sampled at rate *p*.

**Figure 2:**
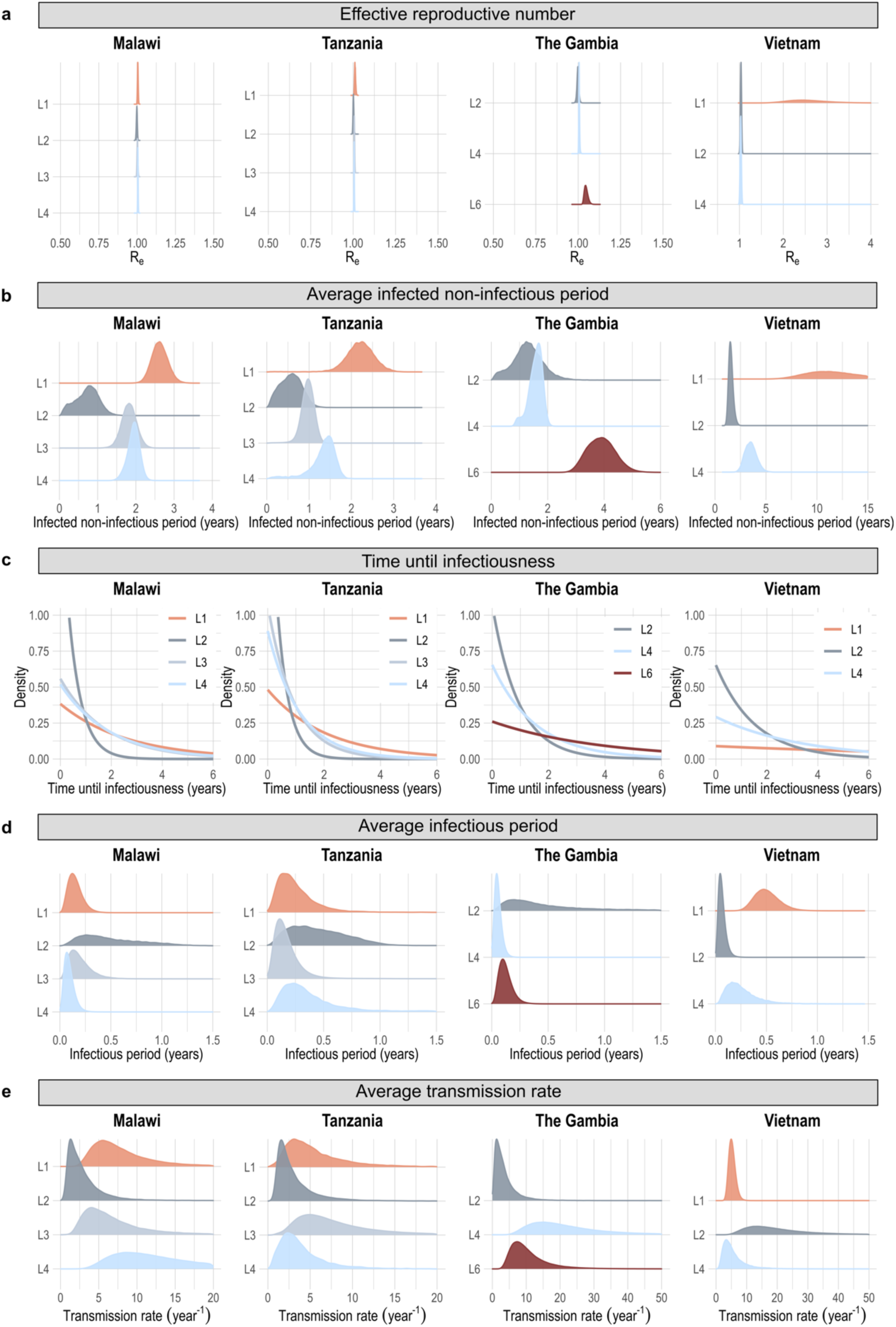
Posterior estimates for the multi-type birth-death model fitted onto genomic data from different sampling locations. **a)** Posterior distributions of the effective reproductive number (R_e_), showing estimates close to 1 for all lineages in all sampling locations (except for L1 in Vietnam). b) Posterior distributions of the average duration of the initial non-infectious period, showing the highest estimates for L1 and L6 in all sampling locations. c) Population distributions of the time until infectiousness, assuming an exponential distribution with rate parameter corresponding to the posterior mean in b). d) Posterior distributions of the average duration of the infectious period, showing no significant differences between lineages. e) Posterior distributions of the average transmission rate during the infectious period, showing no significant differences between lineages.

**Table 1:** Posterior mean and 95% highest posterior density interval for the estimated proportion of infected individuals who progress to infectious TB within one year of infection.

|  | Lineage 1 | Lineage 2 | Lineage 3 | Lineage 4 | Lineage 6 |
| --- | --- | --- | --- | --- | --- |
| Malawi | 0.32 [0.28,0.36] | 0.86 [0.46,0.99] | 0.43 [0.36,0.49] | 0.41 [0.36,0.46] | - |
| Tanzania | 0.38 [0.29,0.46] | 0.91 [0.52,1.00] | 0.66 [0.54,0.75] | 0.59 [0.41,0.78] | - |
| The Gambia | - | 0.66 [0.25,0.93] | - | 0.48 [0.38,0.60] | 0.23 [0.18,0.28] |
| Vietnam | 0.086 [0.049,0.12] | 0.48 [0.39,0.56] | - | 0.25 [0.19,0.32] | - |

To assess the robustness of our results to prior assumptions, we repeated the analyses assuming different levels of underreporting (see Materials and Methods). This resulted in different absolute values for the posterior estimates, but similar relative differences between lineages (Figure S1, Figure S2). Moreover, we also allowed for some rates to change 30 years before the most recent sample, to rule out the influence of independent MTBC introduction events that potentially shape the early parts of the phylogenetic trees (see Materials and Methods). To limit the model complexity for these analyses, we fixed R_e_ to 1 for both time intervals. The parameter estimates for the most recent time interval (Figure S3) were similar to the estimates from the main analyses (Figure 2), suggesting limited biases due to independent introductions. Finally, since the priors in the main analyses put an infinitely small weight on very short non-infectious periods (Table S1), we reparametrized the model and tested the effect of a stronger prior support for short non-infectious periods (see Materials and Methods). Again, the relative differences between lineages remained unchanged (Figure S4).

We further investigated the importance of the non-infectious period in explaining the genomic data by comparing the multi-type birth-death estimates to the estimates from the single-type birth-death model, where infected individuals are assumed to instantaneously become infectious (Figure 1b; Table S2). For the single-type birth-death model, the estimated R_e_ was again close to 1 for all lineages at all locations under study, except for L1 in Vietnam (Figure S5; Table 2). The estimates for the expected time until secondary transmission were also similar for both models, suggesting the longest times for L1 and L6 (Figure S6; Table 2). This average time until transmission is related to commonly used transmission metrics like clustering rates and TBLs. Similarly, the estimates for the average total infected period were in good accordance, indicating robustness to the choice of model (Figure S7; Table 2). As expected, the major difference between the two models was that the estimates from the single-type birth-death model suggest a relatively long infectious period associated with a relatively low transmission rate, while the results from the multi-type birth-death model suggest a non-negligible non-infectious period, followed by a relatively short infectious period associated with a high transmission rate (Figure S8; Figure S9; Table 2). To investigate which of these two models best explains the data, we performed a model selection analysis with 50% prior weight on each model (see Materials and Methods). This analysis resulted in a clear posterior preference (83-100% posterior support) for the multi-type birth-death model for all lineages except L2 (Table 3), which was not surprising given the consistently short non-infectious period estimated for this lineage (Figure 2b). Together, these results further confirm that the longer time between transmission events for “ancient” lineages, as observed in this and previous studies, stems from a later onset, rather than a lower level of infectiousness.

**Table 2:** Arithmetic expressions used to calculate the posterior distributions of parameters that were not directly part of the model.

| Parameter | Multi-type birth-death model | Single-type birth-death model |
| --- | --- | --- |
| Transmission rate during infectiousness | $R_{e,M}\delta_M$ | $R_{e,S}\delta_S$ |
| Time between start of infection and first transmission event | $1/\sigma + 1/\lambda_M$ | $1/\lambda_S$ |
| Non-infectious period | $1/\sigma$ | 0 |
| Infectious period | $1/\delta_M$ | $1/\delta_S$ |
| Total infected period | $1/\sigma + 1/\delta_M$ | $1/\delta_S$ |

**Table 3:** Posterior probability of the multi-type birth-death model, assessed in a model selection analysis assuming 50% prior probability of the single-type (0) and multi-type (1) birth-death model.

|  | Lineage 1 | Lineage 2 | Lineage 3 | Lineage 4 | Lineage 6 |
| --- | --- | --- | --- | --- | --- |
| Malawi | 1 | 0.48 | 1 | 1 | - |
| Tanzania | 0.98 | 0.35 | 0.95 | 0.83 | - |
| The Gambia | - | 0.72 | - | 1 | 1 |
| Vietnam | 1 | 1 | - | 1 | - |

The branch lengths in the phylogenetic tree, informing our parameter estimates, do not only depend on the actual time between transmission events, but also on the clock rate (number of substitutions per site per year) which is used to convert the observed genetic changes into time. In our initial analyses, we estimated the clock rate from the data, setting a relatively informative prior and allowing for lineage-specific clock rate estimates (see Materials and Methods; Table S3). However, estimates for the clock rate and time between transmissions might be confounded due to the limited clock signal in the data. To investigate the impact of this on our results, we fixed the lineage-specific clock rates to a set of values and examined the order of magnitude of difference required to explain the observed differences in branch lengths, from which the differences in the non-infectious period duration are derived. We focused on L1 and L2 in Tanzania, as these showed strong differences in the estimated non-infectious period (Figure 2b). The results show that the L1 clock rate would need to be 16 to 64-fold higher than the L2 clock rate in order for the posterior distributions of these estimates to overlap (Figure 3). This estimated difference is much larger than the two-fold difference reported before (Menardo et al. 2019) and estimated in our main analyses (Table S3). This suggests that while assumptions about the clock rate do have a significant impact on epidemiological parameter estimates, the expected clock rate differences between lineages are not sufficient to explain the observed differences in branch lengths.

**Figure 3:**
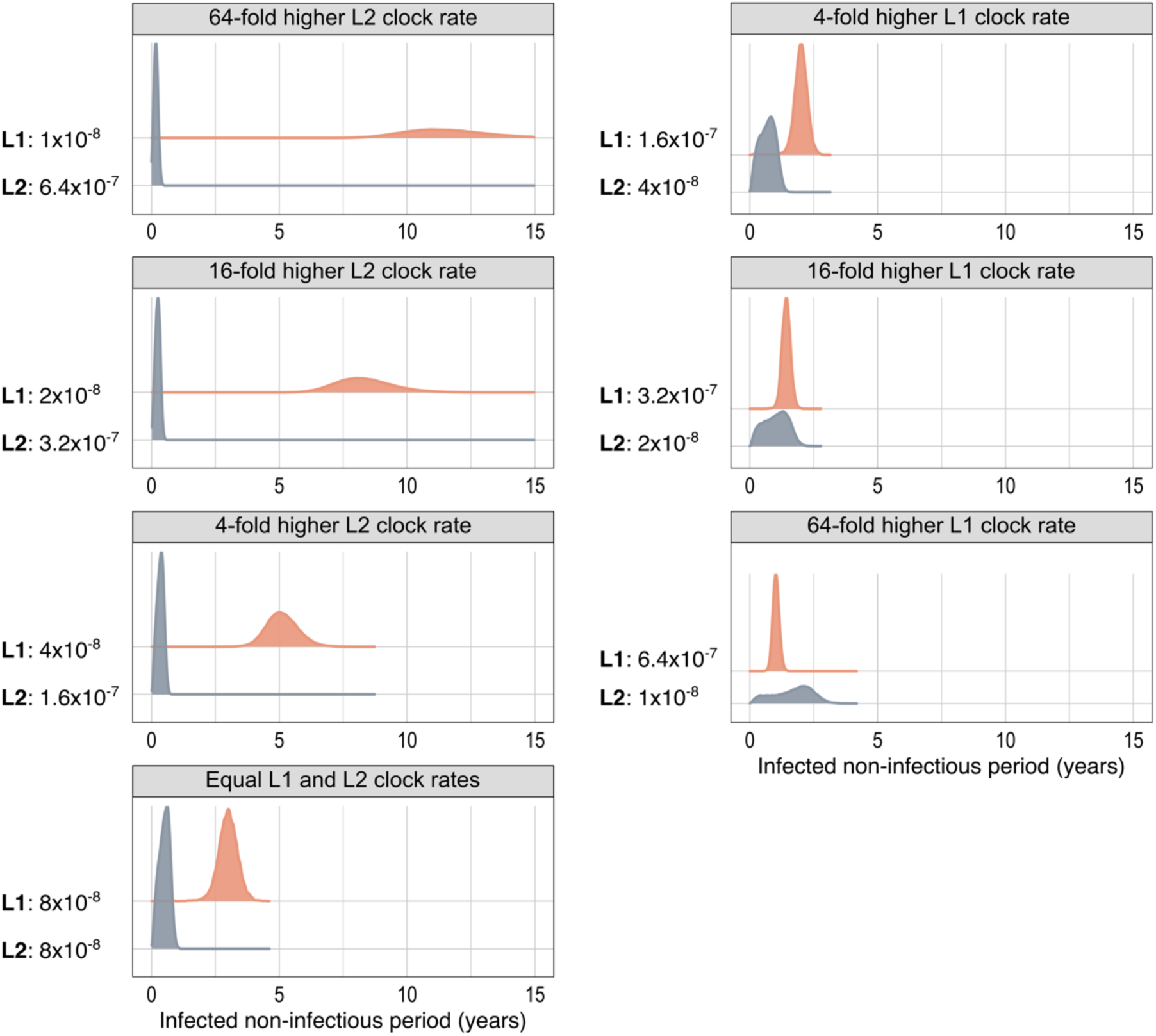
Posterior estimates of the non-infectious period under the multi-type birth-death model, using different lineage-specific clock rates. Posterior distributions of the average duration of the initial non-infectious period for L1 and L2 in Tanzania, assuming different combinations of fixed clock rates (substitutions/site/year). The distributions start overlapping when the clock rate for L1 is at least 16-fold higher than for L2.

## Discussion

In this study, we applied phylodynamic birth-death models to genomic data collected in four different locations where “ancient” and “modern” MTBC lineages co-circulate, with the aim of inferring lineage-specific epidemiological characteristics. Previous studies have suggested a lower transmissibility of “ancient” lineages (mainly L1) compared to “modern” lineages (mainly L2), based on clustering rates or TBLs (Yang et al. 2012; Guerra-Assunção et al. 2015; Holt et al. 2018; Sobkowiak et al. 2020; Freschi et al. 2021; Zwyer et al. 2023; Gröschel et al. 2024), both of which indirectly measure the time between subsequent transmission events. In accordance with these previous findings, our results support a longer expected time between transmission events for L1 and L6 compared to L2, L3, and L4. By explicitly modelling (non-)infectiousness in infected individuals, we show that this difference can be robustly explained by a longer initial period of non-infectiousness in patients infected with L1 or L6 strains. Our results further show that this non-infectious period is followed by an infectious period for which the average duration and level of infectiousness (measured as the transmission rate, i.e. the rate of secondary cases generation per infected individual) is not significantly different between lineages. We demonstrate that a longer initial non-infectious period for “ancient” lineages better explains the data than the alternative model of a longer infectious period starting immediately upon infection, combined with a lower transmission rate (resulting from a lower level of infectiousness) during this infectious period. Overall, our findings are in accordance with a previous household contact study showing that strains from L4 and L6 are associated with similar levels of patient infectiousness, but that L4 is associated with an increased risk of disease progression within 2 years of infection (de Jong et al. 2008).

The observed association between “ancient” lineages and slow disease progression could be linked to the stronger inflammatory immune response reported against these strains (Portevin et al. 2011; Chen et al. 2014) as well as their lower replication rate (Reiling et al. 2013; Sanoussi et al. 2017; Hiza et al. 2024), both of which have been suggested to reflect lower virulence. Another manifestation of this reduced virulence is the observed association of L1 and L6 with old age (Thwaites et al. 2008; de Jong et al. 2010; Guerra-Assunção et al. 2015) as well as with HIV co-infection (Glynn et al. 2010; de Jong et al. 2010; Guerra-Assunção et al. 2015; Asante-Poku et al. 2016), suggesting that L1 and L6 might be more likely to cause asymptomatic, non-infectious infection in individuals that are not immunosuppressed. In support of this notion, the average asymptomatic period was estimated to be longer in countries with a high L1 burden (Ku et al. 2021), and a higher prevalence of subclinical TB has recently been observed in patients infected with “ancient” lineage strains (Long et al. 2024).

Our estimates of the onset of infectiousness are population averages, summarizing an underlying population distribution. Assuming that this population distribution is exponential (Borgdorff et al. 2011; Behr et al. 2018; Menzies et al. 2021) and that 75% of all cases are not reported, our estimates suggest that between 23% (L6) and 73% (L2) of the population progresses to infectious TB within the first year of infection. These values are within the range of previous estimates (Borgdorff et al. 2011; Sloot et al. 2014; Behr et al. 2018; Emery et al. 2021; Menzies et al. 2021; Horton et al. 2023), although there is variation across countries. Assuming lower rates of underreporting resulted in lower estimated rates of disease progression. However, the exact level of underreporting is unknown and likely varies across sampling locations.

The average time between transmission events is often used as a measure for transmissibility but is not the only factor determining how rapidly the prevalence of MTBC lineages changes over time. Instead, these dynamics are also largely determined by the average number of secondary cases caused by one infected individual (R_e_), which is the product of the average duration of infectiousness and the transmission rate during infectiousness. Our results show that the average R_e_ is consistently estimated around one for all lineages in all locations under study. The exception to this is L1 in Vietnam, which is potentially due to the dominance of sublineage L1.1.1, reported to demonstrate increased transmission potential (Stanley et al. 2024), whereas L1.1.2, L1.1.3, L1.2.1, and L1.2.1 are circulating in Malawi and Tanzania. Although our R_e_ estimates represent time averages, and we did not investigate changes in R_e_ through time, an R_e_ of one implies that the TB prevalence per lineage remains relatively constant over time. However, the longer period between transmission events observed for L1 and L6 implies a lower turnover rate within the population of infectious individuals.

Genetic distances between sampled isolates, and consequently also branch lengths in a phylogenetic tree, are not only determined by the rate of transmission, but also by the clock rate and the sampling density. In contrast to clustering methods (applying a fixed SNP threshold) and methods based on TBLs, which both implicitly assume equal clock rates for all lineages, the phylodynamic models used here allow for the simultaneous inference of clock rates from the genomic data. Since the clock signal in *M. tuberculosis* data is intrinsically weak, we tested different scenarios by fixing the clock rate to a set of different values. These analyses show that, in order for the clock rate only to explain the branch length differences, the clock rate for L1 would need to be 16 to 64-fold higher than for L2, which is considerably more than the two-fold difference estimated in a previous systematic analysis (Menardo et al. 2019).

It is worth noting that the non-infectious phase at the start of infection could be associated with reduced bacterial replication and, consequently, a lower mutation rate. While there is some evidence for mutagenesis occurring during the non-infectious phase (Ford et al. 2011; Lillebaek et al. 2016; Colangeli et al. 2020), a reduced mutation rate could bias the estimated duration of this phase, especially when only the genetic diversity is taken into account (as is done in clustering and TBL analyses). In our phylodynamic analyses, we cannot assign different rates to the different infectious stages but assume an average rate. Future methodological work on assigning such different rates will enable the quantification of potential differences.

Except for the L6 culture bias (Sanoussi et al. 2017) accounted for in the priors, we assumed no lineage-specific sampling biases. Nonetheless, the datasets used in this study mainly contain isolates from patients who presented themselves at a healthcare center, most likely following symptom development. While this includes patients who potentially went through an asymptomatic/subclinical phase before progressing to active TB disease, it does not include infected individuals who never develop symptoms. The incidence of subclinical cases who never get diagnosed is currently unknown but might be higher in individuals infected with “ancient” lineage strains (Long et al. 2024). If this is indeed the case, this would imply a lineage-specific undersampling and might affect our results. Without genomic data collected from these fully asymptomatic cases, such biases are challenging to control for. However, these cases are only relevant for the transmission dynamics if they do not represent dead ends in transmission chains, which would imply that they are infectious despite being asymptomatic. While some studies do suggest some degree of infectiousness in subclinical cases (Xu et al. 2019; Frascella et al. 2021; Lau et al. 2022), more studies are needed to determine the strength and variation in infectiousness in these individuals.

Except for the differential presence of the TbD1 genomic region (Brosch et al. 2002) and some recently identified regions of difference (Behruznia et al.), little is known about the genetic differences between “ancient” and “modern” lineages that may underlie the differential TB progression rate observed in this study. Furthermore, the relevance of the “ancient/modern” dichotomy can be questioned for several reasons. First, this and previous studies show clear epidemiological differences between L1/L6 and L2, but L3 and L4 seem to show intermediate behavior. Second, this classification does not properly account for the recently discovered lineages L7-10. Additionally, within-lineage diversity, especially within the most diverse L1 (Coscolla and Gagneux 2014), might further complicate the picture. This calls for a more fine-grained epidemiological characterization, which would require an unbiased sample set representing the genetic diversity in the MTBC at higher resolution.

Taken together, our results demonstrate that the MTBC lineages circulating in Malawi, Tanzania, The Gambia, and Vietnam are associated with a similar effective reproductive number, but different onset of infectiousness. In particular, the slower progression to an infectious disease state, as observed for L1 and L6, results in longer times between transmission events. These results can explain why the prevalence per lineage tends to stay relatively stable over time, despite the higher incidence of “modern” compared to “ancient” lineages in these settings. Our findings provide insights into the epidemiological consequences of MTBC genetic diversity, but more studies are needed to narrow down the underlying genetic determinants.

## Materials and Methods

### Study populations

#### Malawi

We retrieved publicly available sequences of isolates collected from adults with culture-confirmed pulmonary or extrapulmonary TB diagnosed through passive case finding at the hospital and peripheral health centers in Karonga District, northern Malawi, between 1995 and 2011 (*n*=1,684) (raw reads available in the European Nucleotide Archive under project accession numbers PRJEB2358 and PRJEB2794) (Guerra-Assunção et al. 2015). The lineage distribution in the genomic dataset is as follows: L1: *n*=266, L2: *n*=70, L3: *n*=188, L4: *n*=1160 (subsampled to *n*=400 for computational feasibility). The reported incidence of smear-positive TB in adults in the district during the sampling period corresponds to 87-124 cases per 100,000 people per year (Guerra-Assunção et al. 2015).

#### Tanzania

We used previously sequenced isolates (*n*=921) from a cohort of sputum smear-positive and GeneXpert-positive adult pulmonary TB patients. These patients were prospectively recruited at the Temeke District hospital in Dar es Salaam, Tanzania, between 2013 and 2019, in the context of the National TB and Leprosy Programme - Tanzania (raw reads available in the European Nucleotide Archive under project accession number PRJEB49562) (Zwyer et al. 2023). The lineage distribution in the genomic dataset is as follows: L1: *n*=137, L2: *n*=74, L3: *n*=426 (subsampled to *n*=400), L4: *n*=284. In 2020, 3,994 TB cases were notified in Temeke (Jerry Hella, personal communication).

#### The Gambia

We used previously sequenced isolates (*n*=1,086) collected in the context of a cluster randomized trial (ClinicalTrials.gov Identifier: NCT01660646) conducted between 2012 and 2014 in the Greater Banjul Area, The Gambia (raw reads available in the European Nucleotide Archive under project accession number PRJEB53138). Patients in the control arm were diagnosed through passive case finding, whereas patients in the intervention arm were diagnosed through a combination of passive and enhanced case finding, although no impact of the intervention on the transmission dynamics was observed (Gehre et al.). The lineage distribution in the genomic dataset is as follows: L2: *n*=35, L4: *n*=735 (subsampled to *n*=400), L6: *n*=316. The reported TB incidence in The Gambia during the sampling period corresponds to 176 per 100,000 people per year (World Bank 2024).

#### Vietnam

Raw reads were retrieved from the NCBI BioProject database (accession ID: PRJNA355614). These were obtained from adults with smear-positive pulmonary TB diagnosed through passive case finding at eight district tuberculosis units in Ho Chi Minh City, Vietnam, between 2008 and 2011 (*n*=1,623) (Holt et al. 2018). The lineage distribution in the genomic dataset is as follows: L1: *n*=380, L2: *n*=1053 (subsampled to *n*=400), L4: *n*=190. The reported annual incidence of pulmonary TB in Ho Chi Minh City during the sampling period corresponds to ∼11,000 cases (Holt et al. 2018). Sampling collection dates were kindly provided by the authors of the study. Since only sampling year and month were available, all dates were set to the 15^th^ of the month.

#### Whole-genome sequence analyses

Whole-genome sequences were analyzed through a variant-calling pipeline developed in house. Trimmomatic v0.39 (Bolger et al. 2014) was used to i) remove the Illumina adapters allowing for 2 mismatches, ii) scan the reads with a 5 bp sliding window approach and trim when the median quality per base drops below 20, and iii) discard reads shorter than 20 bp. For paired-end data, SeqPrep v1.3.1 (https://github.com/jstjohn/SeqPrep) was used to identify and merge reads with an overlap of at least 15 bp. The processed reads were aligned to an inferred ancestor of the MTBC (Comas et al. 2010) using BWA mem v0.7.17 (Li and Durbin 2009). Duplicate reads were identified and removed using the MarkDuplicates module of Picard v2.26.2 (http://broadinstitute.github.io/picard/). Sequencing reads were taxonomically classified using Kraken (Wood and Salzberg 2014), and non-*Mtb* mappings were discarded as described previously (Goig et al. 2020). Variant calling was performed using the mutect2 module of GATK v4.2.4.1 (McKenna et al. 2010). Variants were then filtered using the FilterMutectCalls in microbial mode. Supplementary and secondary alignments were excluded (Mariner-Llicer et al.), as well as genomic positions in repetitive regions such as PE, PPE, and PGRS genes or phages (Stucki et al. 2016). Samples with an average sequencing depth lower than 15X or with more than 1% of contaminating reads from non-tuberculous mycobacteria were excluded from downstream analysis. Lineages were identified based on SNPs as described in Coll et al. (2014).

#### Multiple sequence alignments

The VCF of all positions was used to create a consensus fasta sequence per isolate. Chromosomal positions that were covered by less than 7 reads, as well as unfixed positions (variant frequency between 10 and 90%), were treated as missing data. An alignment of polymorphic positions was generated for all sequences per lineage per location, by concatenating all high-quality SNPs, excluding sites that had more than 10% of missing data, as well as drug-resistance-related sites and repetitive regions.

#### Multi-type birth-death model

We fit a multi-type birth-death model to the sequence alignments (Kühnert et al. 2016) (Figure 1a), where non-infectious and infectious individuals are treated as two different host types. Under this model, only infectious individuals can transmit (occurring at rate λ_M_), resulting in a new individual in the ‘infected non-infectious’ compartment. Individuals in this compartment cannot transmit, but migrate to the infectious compartment at a constant rate σ (we assume no back migration). Infectious individuals become non-infectious due to recovery, death or removal through sampling, occurring at a constant rate δ_M_, implying that they get removed from the system. This compartmental setup effectively implies that all individuals who eventually progress to infectious disease first go through a phase of non-infectiousness. Hence, infected individuals who never become infectious are not part of the system under study. Instead of λ_M_, the model was parametrized with *R_e,M_* (which equals λ_M_/δ_M_) as more prior knowledge is available for this parameter (Loiseau et al. 2023; Zwyer et al. 2023; Windels et al. 2024). Infected individuals are sampled with sampling proportion *p*, which was set to zero before the onset of sampling and set to a fixed, non-zero value afterwards. This value was calculated per sampling location as the total number of sequences divided by the number of cases reported during the sampling period and multiplied by 0.25 to reflect 75% underreporting of TB cases (this underreporting level was varied in the sensitivity analyses; see below). For L6, we took a culture bias into account by assuming that the efficiency of culture growth for L6 is two third of that for L4 (Sanoussi et al. 2017). Upon sampling an infectious patient, the patient was assumed to be removed from the infectious pool with probability *r* (Gavryushkina et al. 2014). We further assumed a strict molecular clock and a general time-reversible nucleotide substitution model with four gamma rate categories to account for site-to-site rate heterogeneity (GTR+ρ_4_). All parameters and their prior distributions are listed in Table S1.

#### Single-type birth-death model

The single-type birth-death model (Stadler 2010) (Figure 1b) represents a constrained version of the multi-type birth-death model, assuming that individuals instantaneously become infectious upon infection. Hence, all infected individuals are infectious and transmit at a constant rate λ_S_. Individuals become non-infectious through recovery, death or removal through sampling at a constant rate 8_S_. Instead of λ_S_, the model was parametrized with *R_e,S_*, which equals λ_S_/8_S_. All other elements of the model, including the sampling, clock, and nucleotide substitution models, are the same as in the multi-type birth-death model. All parameters and their prior distributions are listed in Table S2.

#### Phylodynamic inference

We performed phylodynamic inference using the bdmm package (Kühnert et al. 2016) v1.0.3 (https://github.com/tgvaughan/bdmm/releases/tag/v1.0.3-unofficial), feast package v8.3.1 (https://github.com/tgvaughan/feast/releases/tag/v8.3.1), and skylinetools package v0.2.0 (https://github.com/laduplessis/skylinetools/releases/tag/0.2.0) in BEAST v2.6.6 (Bouckaert et al. 2014; Bouckaert et al. 2019). Data from each lineage in each location were analyzed independently. Variable SNP alignments were augmented with a count of invariant A, C, G, and T nucleotides (Leaché et al. 2015). Alignments containing more than 400 sequences were randomly downsampled to 400 sequences for computational feasibility, and sampling proportions were adjusted accordingly. For each analysis, three independent Markov Chain Monte Carlo chains were run, with states sampled every 1,000 steps and trees sampled every 10,000 steps. Convergence was assessed with Tracer (Rambaut et al. 2018), confirming that the effective sample size (ESS) was at least 200 for the parameters of interest. 10% of each chain was discarded as burn-in, and the remaining samples across the three chains were pooled and downsampled by a factor 10,000 using LogCombiner (Bouckaert et al. 2019), resulting in at least 49,000,000 iterations in combined chains. All phylodynamic inference steps were implemented in a Snakemake workflow (Mölder et al. 2021). Posterior distributions on derived parameters were calculated using the appropriate arithmetic expressions (Table 2). The model assumes that the time until onset of infectiousness is exponentially distributed, with the mean time being the parameter of that exponential distribution. In Figure 2b, Figure S1b, Figure S2b, Figure S3a, and Figure S4b we provide the posterior distribution for this mean time until onset of infectiousness, 1/α. In Figure 2c, Figure S1c, Figure S2c, Figure S3b, and Figure S4c we show the exponential distribution for the most likely mean time parameter, i.e. the distribution of all times until infectiousness onset in the population.

#### Sensitivity analyses

The robustness of the phylodynamic inference to sampling assumptions was assessed by assuming lower levels of underreporting of TB cases (50% and 0%). To investigate the effect of clock rate assumptions, we focused on L1 and L2 in Tanzania, two lineages with largely different estimates for the non-infectious period (Figure 2b). We examined how large the difference in clock rates would need to be, if it were invoked as the only factor underlying the difference in branch lengths. To this end, we fixed the clock rate for each lineage to different values, chosen within the range of previously reported values (Menardo et al. 2019), and checked which combination resulted in similar non-infectious period estimates.

When the evolutionary history of the sampled isolates is characterized by multiple independent introduction events in the sampling locations, the early parts of the phylogenetic trees might not result from the same population dynamic process as the more recent parts. This can lead to a model misspecification and might bias the epidemiological parameter estimates. To eliminate such effects, we repeated the multi-type birth-death analyses, allowing for a change in α and 8_M_ at a point in time set to 30 years before the most recent sample. For computational feasibility, *R_e,M_* was set to 1 for both time intervals.

The Lognormal(0,1) prior distribution on σ in the main multi-type birth-death analyses puts a very small weight on short non-infectious periods. To allow for a non-infectious period close to zero (which would approximately correspond to a single-type birth-death model), we reparametrized the model with 1/σ and tested the effect of an Exp(1) prior (in other words, a prior with high support for low values) on this parameter.

#### Model selection

To perform model selection on the single-type and multi-type birth-death model, we allowed BEAST to select between two models: 1) a model where both σ and the R_e_ between compartments (*R_e,M_*) is zero, while the R_e_ within the infectious compartment (*R_e,S_*) is non-zero (corresponding to the single-type birth-death model), and 2) a model where *R_e,S_* is zero, while σ and *R_e,M_* are non-zero (corresponding to the multi-type birth-death model). To this end, spike and slab priors were set on σ, *R_e,S_* and *R_e,M_*, by defining them as ModelSelectionParameter (feast package) and putting the same priors as before on the non-zero values of σ, *R_e,S_* and *R_e,M_*. An equal prior weight was put on both models.

## Data availability

We used data that are publicly available in the European Nucleotide Archive under the project accession numbers PRJEB2358, PRJEB2794, PRJEB49562, and PRJEB53138, and in the NCBI BioProject database under accession number PRJNA355614. The code for the phylodynamic analyses, including the Snakemake workflow and BEAST2 XML files, is available at https://github.com/EtthelWindels/snakemake-mtb-beast2.

## Acknowledgements

This project has received funding from the European Research Council (ERC) under the European Union’s Horizon 2020 research and innovation programme grant agreement no. 101001077 (to E.M.W. and T.S.) and ETH Zürich (to E.M.W. and T.S.). This work was further supported by the Swiss National Science Foundation (grants CRSII5_213514 and 320030-227432 to S.G.) and the ERC (883582 to S.G.). Calculations were performed on the Euler cluster at ETH Zürich and at sciCORE (https://scicore.unibas.ch/) scientific computing core facility at the University of Basel. We would like to thank Timothy G. Vaughan for help with the phylodynamic analyses, Louis du Plessis for insightful discussions, and Christoph Stritt and Antoine Zwaans for valuable feedback on the manuscript.

## Author contributions

E.M.W., B.d.J., T.S., and S.G. conceived the study. E.M.W., D.B., and C.J.M. performed data curation. C.V.A. and E.M.W. generated the Snakemake workflow. E.M.W. conducted the phylodynamic analyses. C.L., D.B., G.A.G., and M.Z. generated the variant calling pipeline. T.S. and S.G. acquired funding. E.M.W. wrote the manuscript. All authors edited the manuscript.

