## Supplementary Material for "Onset of infectiousness explains differences in transmissibility across *Mycobacterium tuberculosis* lineages"

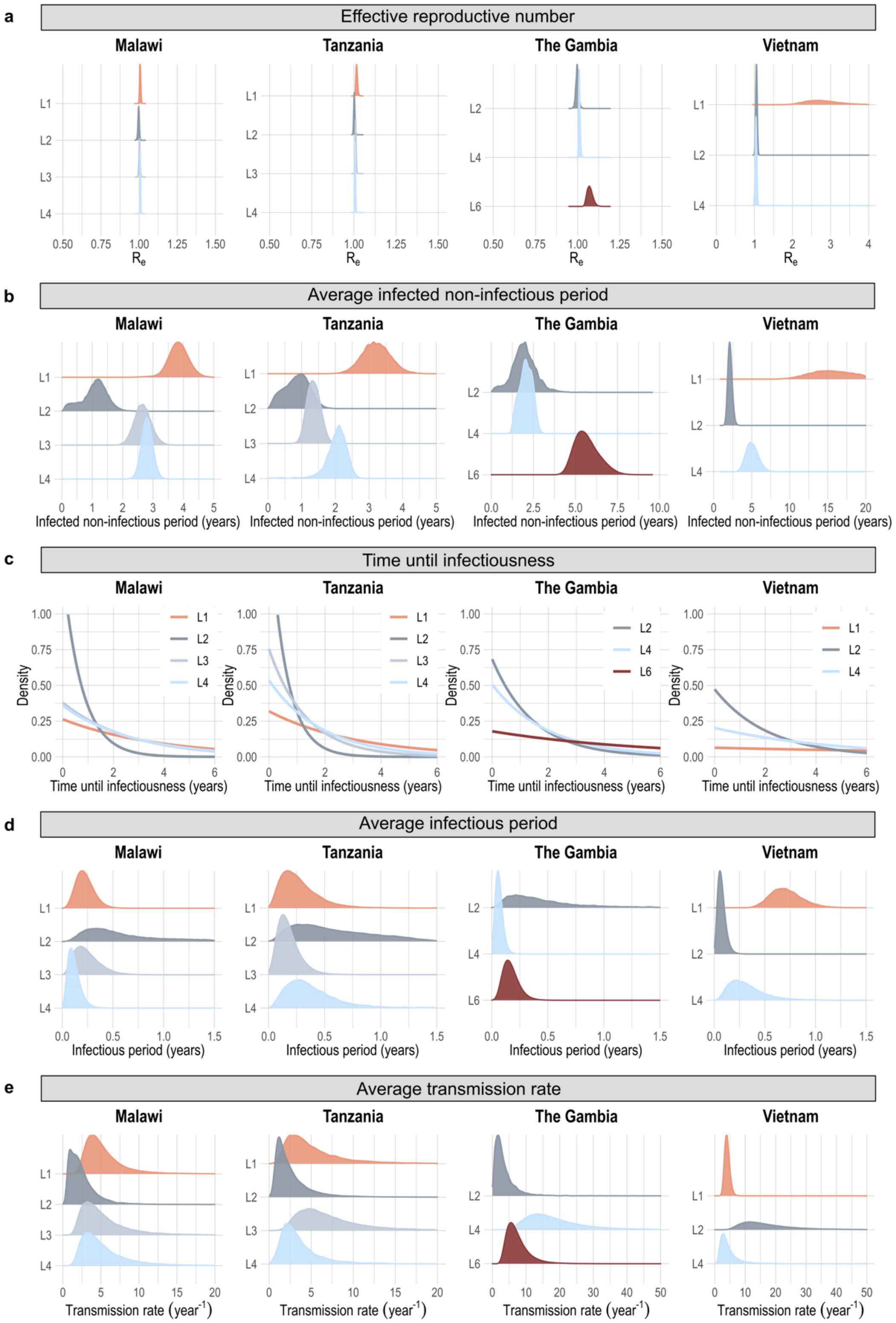


**Figure S1: Posterior estimates for the multi-type birth-death model fitted onto genomic data from different sampling locations, assuming that 50% of the cases were not reported.** a) Posterior distributions of the effective reproductive number ($R_e$). b) Posterior distributions of the average duration of the initial non-infectious period. c) Population distributions of the time until infectiousness, assuming an exponential distribution with rate parameter corresponding to the posterior mean in b). d) Posterior distributions of the average duration of the infectious period. e) Posterior distributions of the average transmission rate during the infectious period.

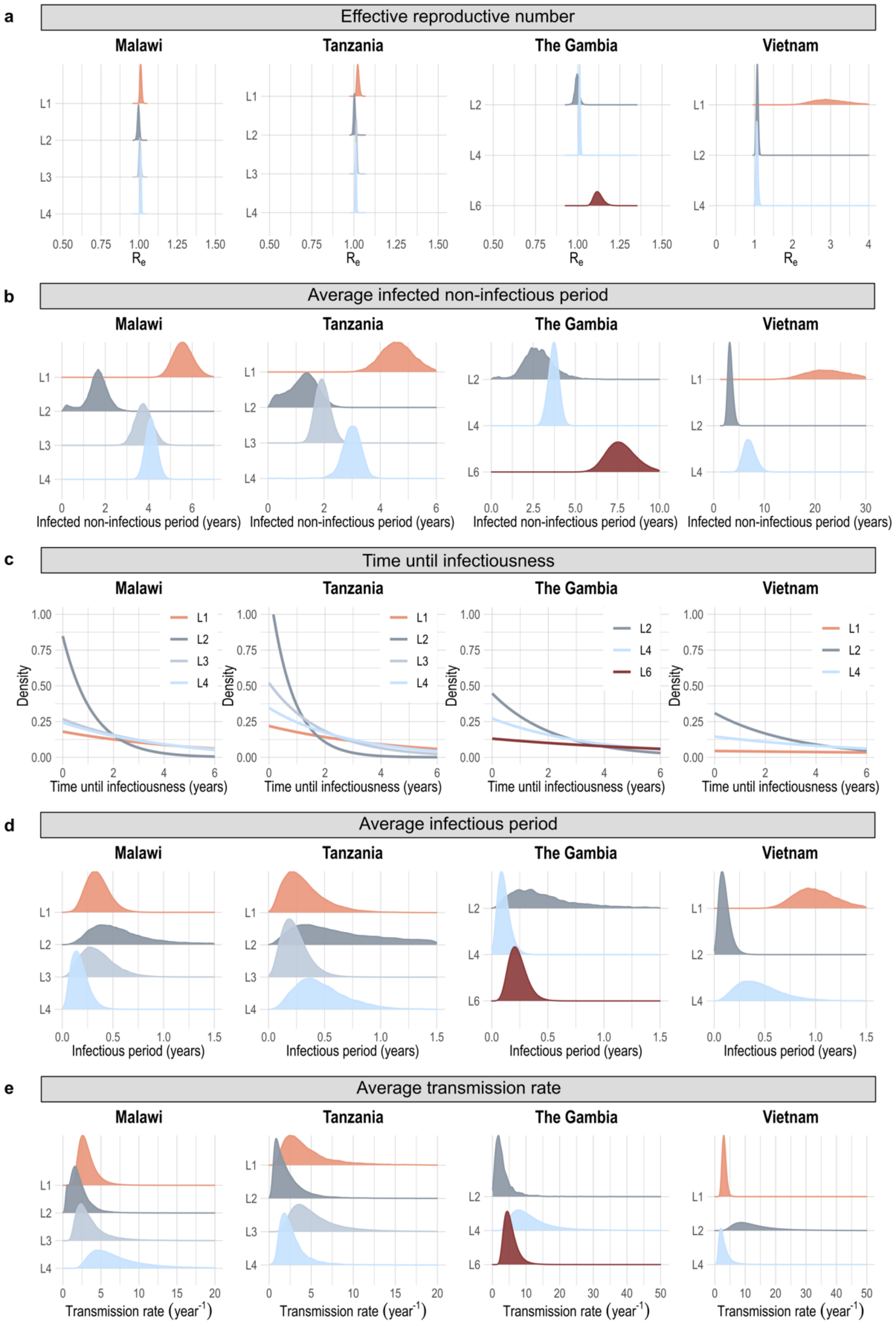


**Figure S2: Posterior estimates for the multi-type birth-death model fitted onto genomic data from different sampling locations, assuming that all cases were reported.** a) Posterior distributions of the effective reproductive number ($R_e$). b) Posterior distributions of the average duration of the initial non-infectious period. c) Population distributions of the time until infectiousness, assuming an exponential distribution with rate parameter corresponding to the posterior mean in b). d) Posterior distributions of the average duration of the infectious period. e) Posterior distributions of the average transmission rate during the infectious period.

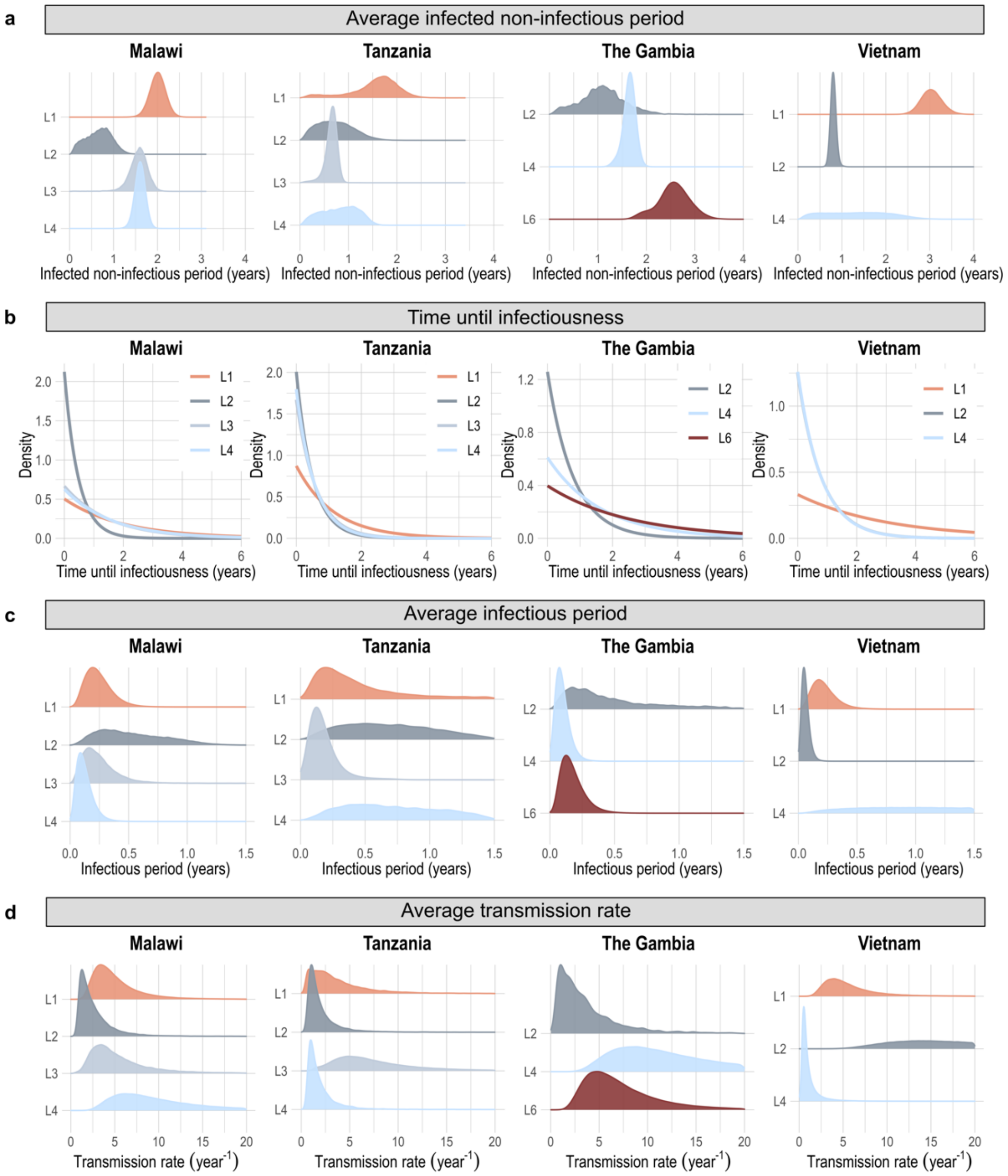


**Figure S3: Posterior estimates for the multi-type birth-death model fitted onto genomic data from different sampling locations, with time-dependent estimates of the non-infectious and infectious periods.** The estimates of $\sigma$ (inverse of the non-infectious period) and $\delta_M$ (inverse of the infectious period) were allowed to change 30 years before the most recent sample, to filter out potential effects of multiple MTBC introductions into the study locations. The $R_e$ was fixed to 1 for both time intervals. Only the estimates for the most recent time interval are shown. a) Posterior distributions of the average duration of the initial non-infectious period. b) Population distributions of the time until infectiousness, assuming an exponential distribution with rate parameter corresponding to the posterior mean in a). c) Posterior distributions of the average duration of the infectious period. d) Posterior distributions of the average transmission rate during the infectious period.

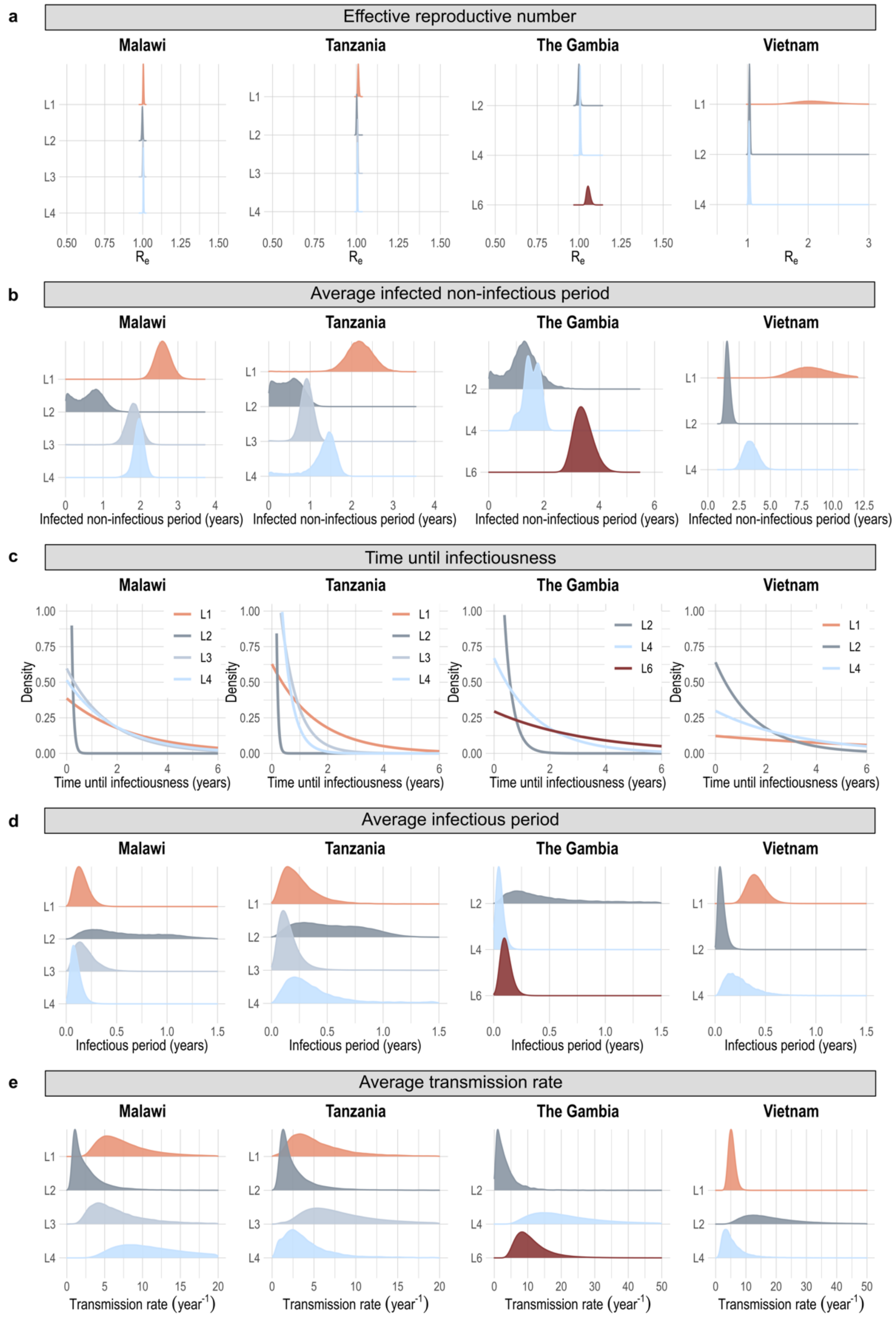


**Figure S4: Posterior estimates for the multi-type birth-death model fitted onto genomic data from different sampling locations, assuming an Exp(1) prior on the non-infectious period.** The model was reparametrized with 1/σ (non-infectious period) instead of σ (rate of becoming infectious). a) Posterior distributions of the effective reproductive number ($R_e$). b) Posterior distributions of the average duration of the initial non-infectious period. c) Population distributions of the time until infectiousness, assuming an exponential distribution with rate corresponding to the posterior mean in b). d) Posterior distributions of the average duration of the infectious period. e) Posterior distributions of the average transmission rate during the infectious period.

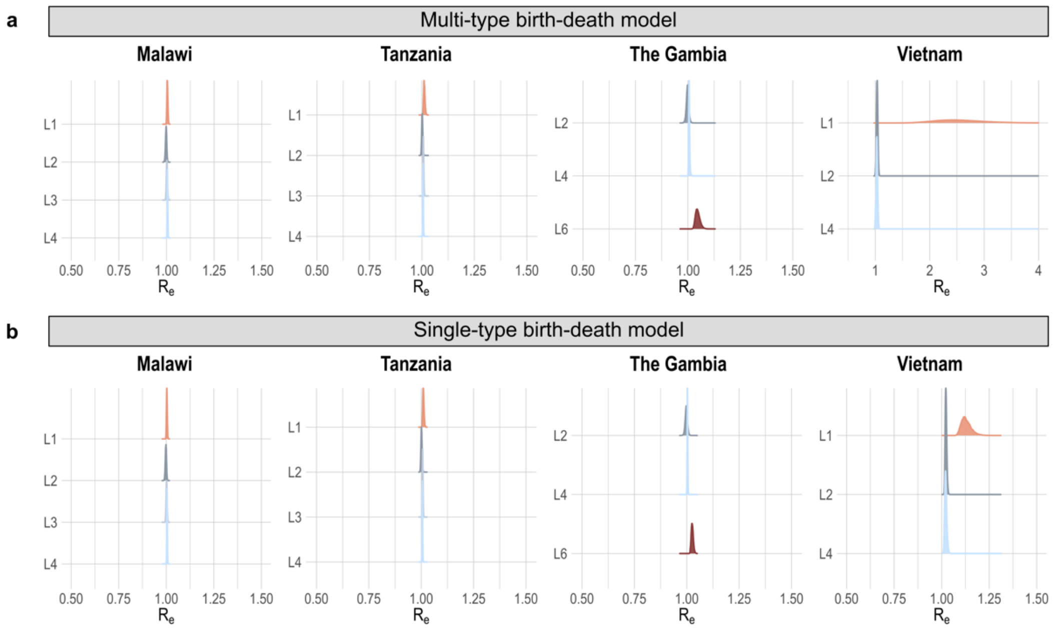


**Figure S5: Comparison of the $R_e$ estimates under a multi-type and single-type birth-death model.** a) Posterior distributions of the effective reproductive number ($R_e$) estimates under a multi-type birth-death model and corresponding to $\lambda_M/\delta_M$. b) Posterior distributions of the $R_e$ estimates under a single-type birth-death model and corresponding to $\lambda_S/\delta_S$. See main text for details on the parameters.

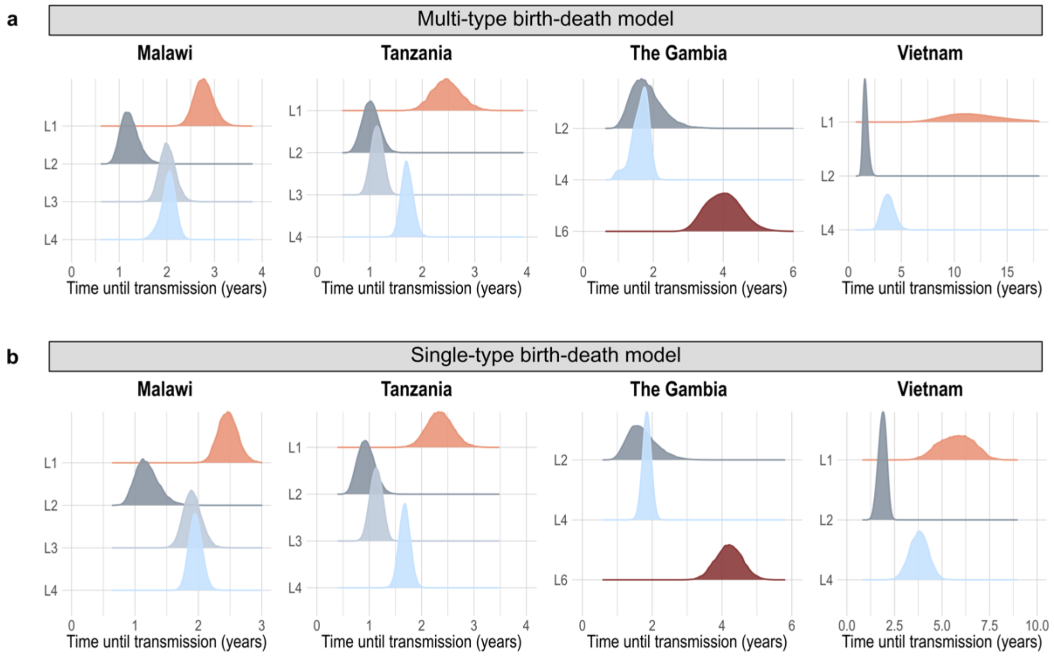


**Figure S6: Comparison of the time until transmission estimates under a multi-type and single-type birth-death model.** a) Posterior distributions of the average time until secondary infection, estimated under a multi-type birth-death model and corresponding to $1/\sigma+1/\lambda_M$. b) Posterior distributions of the average time until secondary infection, estimated under a single-type birth-death model and corresponding to $1/\lambda_S$. See main text for details on the parameters.

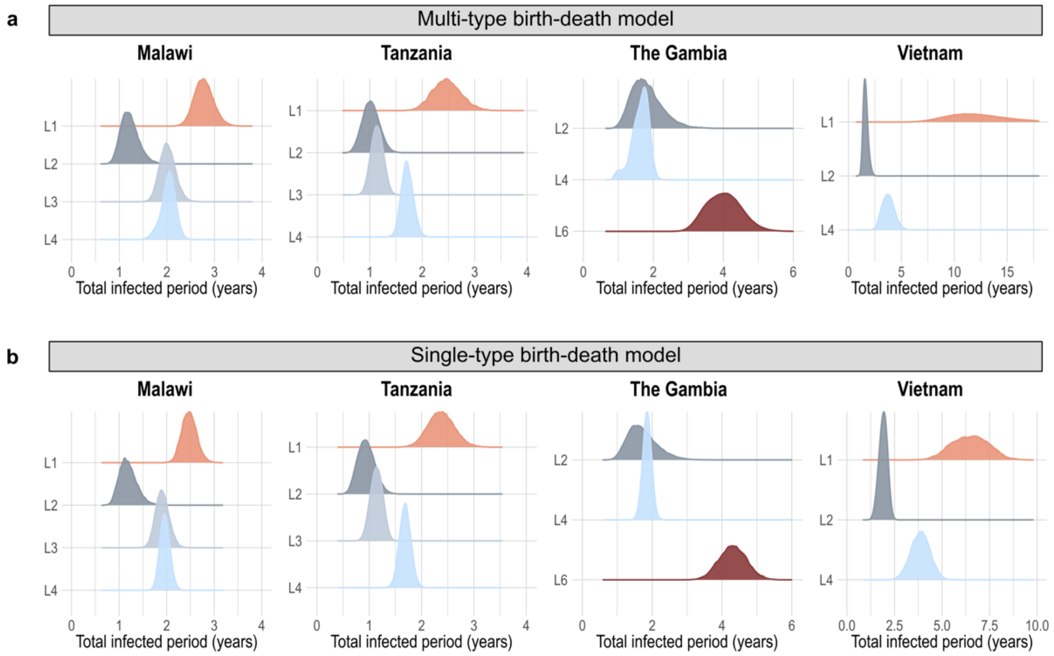


**Figure S7: Comparison of the total duration of infection estimates under a multi-type and single-type birth-death model.** a) Posterior distributions of the average total duration of infection, estimated under a multi-type birth-death model and corresponding to $1/\sigma+1/\delta_M$. b) Posterior distributions of the average total duration of infection, estimated under a single-type birth-death model and corresponding to $1/\delta_S$. See main text for details on the parameters.

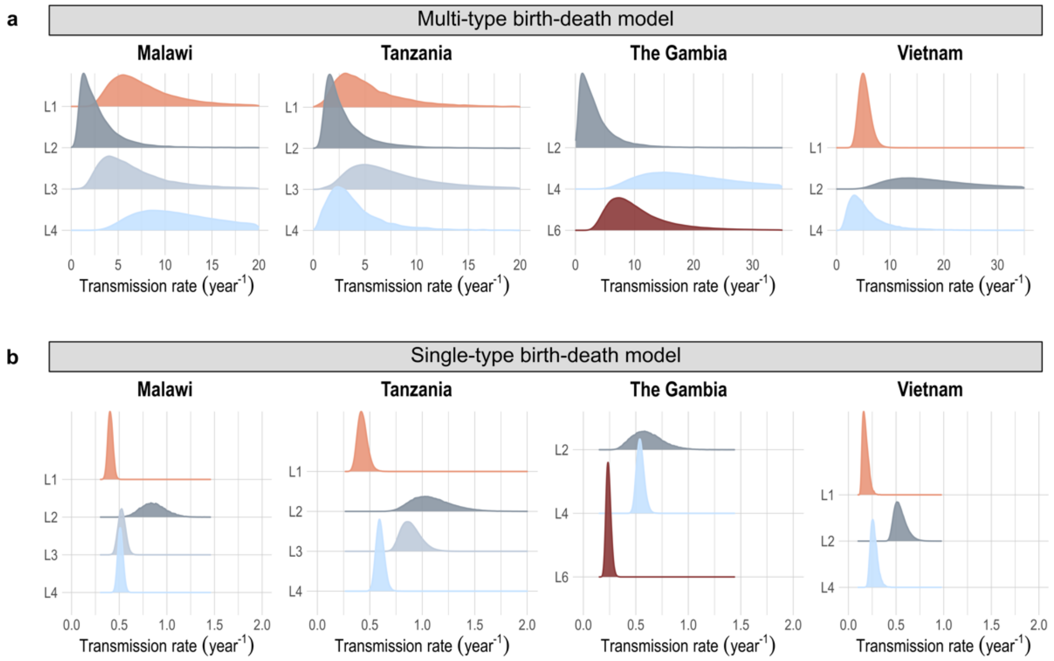


**Figure S8: Comparison of the transmission rate estimates under a multi-type and single-type birth-death model.** a) Posterior distributions of the average transmission rate, estimated under a multi-type birth-death model and corresponding to $\lambda_M$. b) Posterior distributions of the average transmission rate, estimated under a single-type birth-death model and corresponding to $\lambda_S$. See main text for details on the parameters.

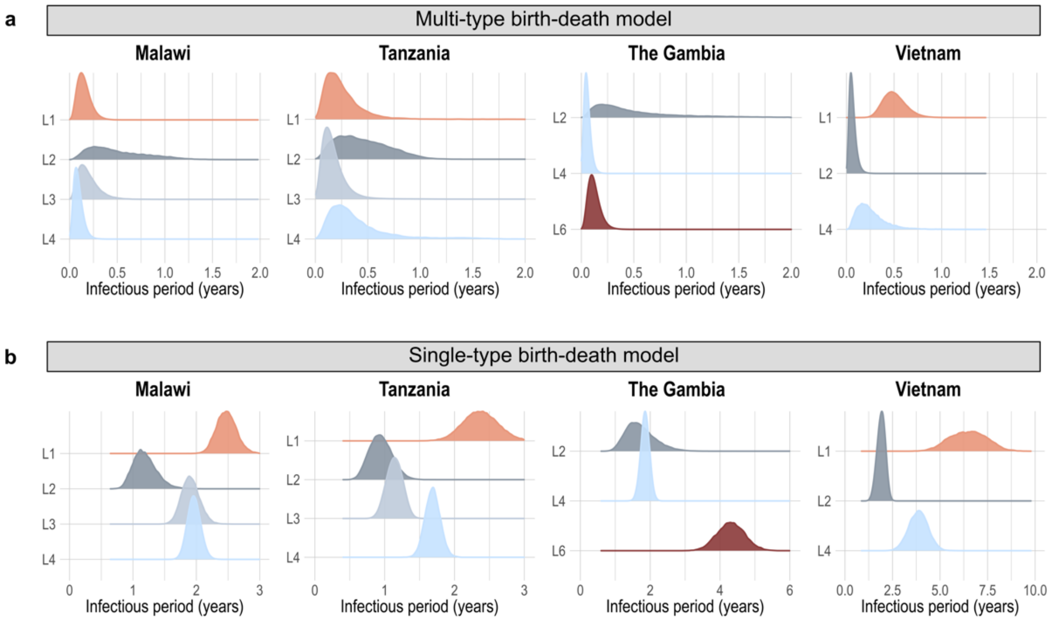


**Figure S9: Comparison of the infectious period estimates under a multi-type and single-type birth-death model.** a) Posterior distributions of the average infectious period, estimated under a multi-type birth-death model and corresponding to $1/\delta_M$. b) Posterior distributions of the average infectious period, estimated under a single-type birth-death model and corresponding to $1/\delta_S$.

**Table S1:** Prior distributions for the parameters of the multi-type birth-death model (see Materials and Methods for more information about the individual parameters)

| **Parameter** | **Symbol** | **Prior** |
|---|---|---|
| Effective reproductive number | $R_{e,M}$ | Lognormal(0,1) |
| Rate of becoming uninfectious | $\delta_M$ | Lognormal(0,1) |
| Rate of becoming infectious | $\sigma$ | Lognormal(0,1) |
| Sampling proportion | $p$ | $\frac{\#sequences}{\#reported\ cases} * (level\ of\ reporting)$ |
| Level of reporting | | 0.25 |
| Probability of removal upon sampling | $r$ | Uniform(0,1) |
| Clock rate | | Lognormal(-16,1) |
| Time of origin | | Uniform(0,1000) |
| Gamma shape | | Exp(1) |
| A→C substitution rate | | Gamma(0.05,10) |
| A→G substitution rate | | Gamma(0.05,20) |
| A→T substitution rate | | Gamma(0.05,10) |
| C→G substitution rate | | Gamma(0.05,10) |
| G→T substitution rate | | Gamma(0.05,10) |

**Table S2:** Prior distributions for the parameters of the single-type birth-death model (see Materials and Methods for more information about the individual parameters)

| Parameter | Symbol | Prior |
|---|---|---|
| Effective reproductive number | $R_{e,S}$ | Lognormal(0,1) |
| Rate of becoming uninfectious | $\delta_S$ | Lognormal(0,1) |
| Sampling proportion | $p$ | $\frac{\#sequences}{\#reported\ cases} * (level\ of\ reporting)$ |
| Level of reporting | | 0.25 |
| Probability of removal upon sampling | $r$ | Uniform(0,1) |
| Clock rate | | Lognormal(-16,1) |
| Time of origin | | Uniform(0,1000) |
| Gamma shape | | Exp(1) |
| A→C substitution rate | | Gamma(0.05,10) |
| A→G substitution rate | | Gamma(0.05,20) |
| A→T substitution rate | | Gamma(0.05,10) |
| C→G substitution rate | | Gamma(0.05,10) |
| G→T substitution rate | | Gamma(0.05,10) |

**Table S3:** Mean and 95% highest posterior density interval for the clock rate estimates (x$10^{-7}$ substitutions per site per year) resulting from the main phylodynamic analyses

| | **Lineage 1** | **Lineage 2** | **Lineage 3** | **Lineage 4** | **Lineage 6** |
|---|---|---|---|---|---|
| Malawi | 1.05 [0.99,1.10] | 0.68 [0.44,0.90] | 0.68 [0.61,0.76] | 0.78 [0.70,0.96] | - |
| Tanzania | 1.30 [1.05,1.67] | 0.62 [0.39,0.94] | 0.93 [0.66,1.22] | 0.83 [0.76,0.94] | - |
| The Gambia | - | 1.09 [0.42,1.83] | - | 2.01 [0.92,4.76] | 1.64 [1.15,2.31] |
| Vietnam | 5.49 [4.18,6.90] | 1.62 [0.96,2.31] | - | 1.48 [0.84,2.25] | - |
